# Aquamin a marine derived multi mineral attenuates toll like receptor mediated inflammatory responses in macrophages

**DOI:** 10.64898/2026.01.19.700098

**Authors:** Emmanouil Lioudakis, Sakshi Hans, Denise O’Gorman, Shane O’Connell, Margaret Lucitt

## Abstract

Toll-like receptors (TLRs) play a central role in innate immune responses through recognition of pathogen-associated molecular patterns (PAMPs) and damage-associated molecular patterns (DAMPs). Activation of TLRs leads to the induction of inflammatory cytokines via MyD88- and TRIF-dependent signaling pathways. Aquamin, a multi-mineral supplement derived from the marine red algae *Lithothamnion*, is known for its anti-inflammatory properties. In this study, we demonstrate that Aquamin dose-dependently inhibits lipopolysaccharide (LPS)-induced production of pro-inflammatory cytokines (TNF-α, IL-6, IL-1β) and chemokines in both human peripheral blood mononuclear cells (hPBMCs) and murine bone marrow-derived macrophages (mBMDMs), without causing cytotoxicity. Mechanistic studies demonstrate that Aquamin specifically suppressed TRIF dependent IRF3 activation downstream of TLR3 and TLR4. These findings support Aquamin as a promising agent for use as an intervention in counteracting inflammatory diseases states where TLR signalling is implicated.

## 1. Introduction

Toll-like receptors (TLRs) are evolutionarily conserved pattern recognition receptors (PRRs) that play a critical role in detecting pathogens and initiating immune responses [1]. TLRs are expressed on the surface and in intracellular endosomal compartments of various immune cells, including dendritic cells, macrophages, and lymphocytes, as well as nonimmune fibroblasts and epithelial cells (ECs) [2]. TLRs recognize structurally conserved components of microbes known as pathogen-associated molecular patterns (PAMPs), as well as damage-associated molecular patterns (DAMPs) which are endogenous molecules released from damaged or dying cells [3]. Upon recognition, TLRs initiate signalling cascades that trigger both innate and adaptive immune responses, involved in the elimination of pathogens such as bacteria, viruses, fungi, and parasites. To date 10 human and 12 mouse TLRs have been identified with TLR 1-9 conserved across both species [4].

Each TLR recognises specific ligands with Toll-like receptor 4 (TLR4) specifically recognising lipopolysaccharides (LPS), which is a component of the outer membrane of Gram-negative bacteria [5]. In contrast TLR3 detects double-stranded RNA from viruses which can be mimicked using synthetic analogs of double-stranded RNA (dsRNA), such as polyinosinic-polycytidylic acid (Poly(I:C)) [6]. Whereas TLR 9 recognises bacterial CpG DNA sequences and synthetic oligodeoxynucleotides (ODN) containing unmethylated CpG [7]. Upon ligand binding, TLRs recruit two well described adaptor proteins, Myeloid differentiation primary response 88 (MyD88) and TIR domain-containing adaptor inducing interferon -beta (TRIF) [8]. These adaptor molecules initiate further downstream signalling which results in various transcription factor activation that drives the production and secretion of inflammatory cytokines, type I interferons (IFN) and chemokines. All TLRs drive inflammatory responses via MyD88 except for TLR3 [9]. In the case of TLR4 signalling MyD88-dependent signalling triggers a rapid pathway which activates the transcription factor Nuclear factor kappa-light-chain-enhancer of activated B cells (NF-κB) and mitogen-activated protein kinases (MAPKs), promoting early transcription of pro-inflammatory genes (TNF-α, IL-6) and the production of reactive oxygen species (ROS) like nitric oxide (NO) [10]. A second MyD88--independent pathway downstream of endosomal TLR4 trafficking, engages the adaptor molecule TRIF which activates transcription factor IFN regulatory factor (IRF) 3 and type 1 interferon production (IFN-β). This TRIF dependent pathway also contributes to a later phase of sustained NF-κB and MAPK activation [11]. Unlike TLR4, which initiates both MyD88 and TRIF depending signalling events, TLR3 primarily engages with TRIF only [12]. In contrast TLR9 signalling primarily involves recruitment of the adaptor protein MyD88 only [9]. The TLR3 TRIF dependent signalling initiates IRF3 type 1 interferon production as well as alternative cross talk pathways that lead to further IRF3, NF-κB, and MAPK signalling [13]. Both MyD88-NF-κB and the TRIF-IRF3 dependent signalling axes are both central to driving the production of cytokines that coordinate the immune response in cell recruitment, and activation of adaptive immunity [14].

It is now widely recognised, that apart from their central roles in mounting effective immunity against infection, TLR signalling and activation can also be initiated via endogenous sterile mediators contributing to the pathophysiology of chronic inflammatory diseases states including autoimmune disorders and cancer [15]. This sterile inflammation is important for tissue repair and regeneration, but can also lead to the development of various chronic inflammatory diseases [16]. In fibrotic disease states activation of TLR4 on fibroblasts is shown to be a key driver of persistent organ fibrosis and offers a potential therapeutic target for patients with systemic sclerosis [17]. In the gut TLRs are crucial for sensing microbes, maintaining barrier integrity, and initiating immune responses, but their dysregulated activation is known to drive intestinal colitis such as in individuals with TLR gene polymorphisms [18–20]. In patients with inflammatory bowel disease increased expression of TLR4 is observed in epithelial and lamina propria cells, suggesting an important role for TLR4 signalling in inflammation pathology here [21]. This is further supported by TLR4 chronic stimulation in colonic epithelial cells being more susceptible to colonic inflammation, reduced epithelial barrier integrity and driving colitis-associated tumorigenesis [22]. TLR4 expressing epithelial cells having a particular important role in orchestrating the inflammatory colitis microenvironment response here [23]. An interesting study in support of the above effects showed using indigestible carbohydrates present in breast milk inhibit TLR4 providing a protective effect in necrotizing small bowel colitis [24]. Due to their central role in immunity, TLRs are therefore being studied as potential targets not only for vaccine development and therapies for infectious disease but for controlling exacerbated inflammation associated with chronic inflammatory disease states and autoimmune disorders.

Aquamin, is a multi-mineral supplement, derived from the red algae *Lithothamnion* species with evidence of anti-inflammatory effects [25, 26] . It consists of 74 different minerals and is especially rich in calcium and magnesium with calcium and magnesium ratios approximately twelve to one (12:1). Aquamin is sold as a dietary supplement (GRAS 000028; Marigot Ltd., Cork, Ireland) and is used in various products for human consumption in Europe, Asia, Australia, and North America. Aquamin has demonstrated significant improvements in gut microbiome health by enhancing gut microbial diversity and barrier integrity [27] and counteracts pro-inflammatory activity to improve barrier function in gut organoid systems [28]. With results from a more recent clinical study showing that the use of this multi-mineral intervention improves disease-related biomarkers in patients with ulcerative colitis [29] . Studies have further reported that Aquamin supplementation can moderately reduce liver ballooning degeneration and collagen deposition in mice on a high-fat-diet (HFD), with improved metabolic protein profiles [30]. While Aquamin shows strong evidence for the potential to mitigate inflammation via cytokine suppression and provide protection in gut and liver disease states, the detailed cellular mechanistic pathways have not been elucidated.

In this study, we investigate the immunomodulatory effects of Aquamin® on innate immune signalling in macrophage cells. We demonstrate that Aquanim can reduce both cytokine and chemokine responses downstream of TLR4 and TLR3 in a TRIF dependant manner. By dampening TLR4- and TLR3-driven signalling, Aquamin® can modulate innate immune activation and limit excessive inflammatory mediator release. This mechanism provides a biological basis for Aquamin’s reported protective effects in inflammatory diseases studies.

## 2. Materials and Methods

### 2.1 Aquamin®

Aquamin® soluble is a mineral-rich natural product obtained from the calcified fronds of red marine algae of the *Lithothamnion* genus. It is particularly rich in Calcium and Magnesium (ratio approximately 12 : 1) and contains measurable levels of 72 additional trace elements Supplemental Table 1 for full mineral composition of Aquamin^®^ established via an independent laboratory (Advanced Laboratories; Salt Lake City, Utah). The complex mineral profile distinguishes Aquamin® from single-mineral supplements. Aquamin is classified as Generally Recognized As Safe (GRAS) (GRN 000028). It is produced and supplied by Marigot Ltd., Cork, Ireland, and is marketed as a dietary supplement for use in a wide range of food and nutritional products. Aquamin® has also received approval as an Investigational New Drug (IND No. 141600) from the U.S. Food and Drug Administration (FDA), permitting its clinical investigation and evaluation beyond its use as a dietary supplement. All materials used in the studies here, including biological agents, synthetic compounds, and the natural Aquamin product, were confirmed to be endotoxin-free. Microbiological testing results for Aquamin, as documented in its certificate of analysis, are provided in Supplemental Table 2.

### 2.2 Peripheral blood mononuclear cell isolation and culture

Buffy coats from anonymous donors were obtained from The Irish Blood Transfusion Service, National Blood Bank, St. James’s Hospital, James’s Street, Dublin 8. PBMCs were isolated and cultured, following institutional ethical approval. Peripheral blood mononuclear cells (PBMCs) were isolated from buffy coats as previous described [31] using density gradient Histopaque 10771 (Sigma Aldrich, Poole, UK). Cells were resuspended and cultured in complete media (RPMI-1640 supplemented with 10% FBS, l-glutamine (2 mmol/L), penicillin (100 units/mL), and streptomycin (100 µg/mL)) all from Sigma Aldrich, Poole, UK.

### 2.3 Murine bone marrow derived macrophages (BMDMs) isolation and culture

Hind legs were dissected from adult C57BL/6 mice. The epiphyses of each femur and tibia were removed, so that the bone marrow could be accessed from the ends. Bone marrow from the tibiae and femurs were flushed into a 50 ml conical tube with 2-3 ml of complete media (DMEM supplemented with 10% FBS, 2 mM L-Glutamine, 100U/ml penicillin, 100 μg/ml streptomycin (Sigma Aldrich, Poole, UK)) using a 23 Gauge needle. The cell suspension was passed through a 70 μm pore size cell strainer. Bone marrow cells flushed from three individual mice were combined and centrifuged at 200 x g for 5 minutes. Cell pellets were resuspended in 10 ml complete media supplemented with 20 ng/ml recombinant mouse macrophage colony-stimulating factor (rmM-CSF) (Miltenyi Biotec LTD), transferred to T-75 flask and incubated for 72 hours in a humidified 5% CO_2_ cell culture incubator at 37°C. After 72 hours, 5 ml of complete DMEM media supplemented with 20 ng/ml rmM-CSF was added and cells incubated for a further 72 hours before use.

### 2.4 Cell Stimulation

Solutions of Aquamin S (Marigot LTD)) were prepared fresh prior to use in cell culture media depending on cell type. The following toll-like receptor (TLR) agonists were used TLR3 Poly(I:C) HMW (100ug/ml), TLR 9 CpG ODN 1826 (3μM) (both InvivoGen) and TLR4 LPS (100ng/ml) from Escherichia coli O55:B5 (Sigma Aldrich, L6529). hPBMCs or mBMDMs were first treated with Aqaumin (0.5,1,2 mg/ml) for 3h prior to TLR agonist stimulation for a further 4-12h. Following TLR stimulation cell culture media was harvested and stored at -80 °C for ELISA cytokine measurements. Cells were processed for either protein or RNA extraction or fixed for immunofluorescence staining as described below.

### 2.5 Cell viability

The Alamar blue assay was used to access cell viability using Resazurin Sodium Salt (Invitrogen R12204). Cells were plated at 100,000 /well in a 96-well plate and treated with various concentrations of Aquamin S (0.5-2mg/ml dissolved in culture media) and incubated for 12 h. At the end of the incubation period, resazurin solution equal to 10% of the cell culture volume was added to each well and incubation continued in standard culture conditions for a further 4 h at 37°C. The absorbance at 570 nm was read on a BioTek ELx808 using KC Junior software.

### 2.6 Semi quantitative RT-PCR

RNA was extracted from cells using Bioline RNA extraction kit (London, UK) following standard kit protocols. RNA concentration and purity was determined using a NanoDrop 2000 (Life Technologies Ltd., Paisley, UK). RNA samples underwent a DNase 1 (Invitrogen) treatment step, to digest single- and double-stranded DNA. The mRNA sample was then ready to use in reverse transcription reactions for cDNA synthesis using RevertAid Reverse Transcriptase (Thermofisher, EP0442) kit . cDNA was used for targeted gene expression analysed by semi-quantitative RT-PCR using target-specific primers. See Supplemental Table 3 for all primer sequences (Integrated DNA Technologies/Belgium). RT-PCR was carried out using SYBR green GoTaq DNA polymerase (Promega, A6002) on Mx3000P QPCR System (Agilent Technology), 40 cycles @ (95 ^O^C for 15 sec, 62 ^O^C for 1 min, 72 ^O^C for 15 sec) with final hold step at 4 ^O^C. The size of the amplicons were checked using a dissociation curve added to the end of the PCR run for primer sequence specificity. Gene expression fold changes was calculated using the comparative Ct method [2^(-ΔΔCt)] and normalized to a reference housekeeping genes GAPDH or 18S RNA.

### 2.7 SDS-PAGE and western inmmunoblotting

Cell lysis and protein extraction was carried out using RIPA buffer (Trizma Base 50 mM; NaCl 150 mM; EDTA 2 mM, NP-40 0.5% S) supplemented with 1× protease/phosphatase inhibitor cocktail on day of use. The Pierce BCA (Thermo fisher) kit was used to quantity total protein according to the manufacturer’s instructions. Standard western blot techniques were performed as previously described ([31]. Briefly 20 μg total protein was resolved on 12% Tris-Glycine SDS-PAGE gels along with a molecular weight standard (Fisher BioReagents (BP3603-500), EZ-Run™ Pre-stained Protein Ladder, 10-170kDa). The protein resolved gels were transferred using semi-dry gel transfer onto PVDF membrane (Amersham, Buchinghamshire, United Kingdom). Following the transfer, the PVDF membrane was blocked for 2 h with TBS-T containing 5% w/v Marvel and probed with the following primary antibodies, overnight at 4°C. Primary anti mouse rabbit mAb antibodies all at 1:1000 dilution from Cell Signalling Technology : iNOS (13120S), IκBα (4812), phospho-NF-κB p65 (Ser536) ( 3033), NF-κB p65 (8242). This was followed by secondary antibody detection using Anti-rabbit IgG, HRP-linked Antibody (1:5,000) (7074), Cell Signalling Technology. Mouse beta -Actin Antibody (C4) (sc-47778) HRP conjugated, (Santa Cruz Biotechnology, Santa Cruz, CA, US)] at 1:1000 dilution was used as a loading control. Signal detection was performed via enhanced chemiluminescence Pierce™ ECL (Thermo Fisher Scientific) and autoradiography using a Fusion Fx imaging system (Vilber). Densitometry quantification of immune detected bands was carried out using Bio1D software.

### 2.8 Enzyme-linked immunosorbent assay (ELISA)

The concentration of cytokines present in cell culture supernatants was quantified using the following ELISA kits according to manufactures instructions . Human TNF-α (DY210), and IL-6 (DY206) from R & D Systems, Minneapolis, MN, US. Mouse TNF-α, (88-7324-88) and IL-6 (88-7064-88) from Invitrogen. Samples were measured in triplicate, and absorbances read on a BioTek ELx808 using KC Junior software.

### 2.9 Cell Immunofluorescence staining

Cells were fixed in 4% paraformaldehyde (PFA) for 20 min at room temperature and blocked with 2% BSA, 0.3% Triton X-100 for 1 h. The cells were incubated at room temperature for 90 min with primary antibodies and followed by secondary antibody for 60 min, all at room temperature and counterstained with DAPI (4083) nuclear staining. The primary antibodies include, iNOS Rabbit mAb (1:200, 13120S) and IRF-3 Rabbit mAb (1:200, 4302) with secondary antibody detection using Anti-rabbit IgG (Alexa Fluor^®^ 555 Conjugate) (4413) all from Cell Signalling Technology. Cell images were acquired using an inverted light microscope (Zeiss Axiovert 200, Carl Zeiss) and immunofluorescence quantified using Image J software.

### 2.10 Statistical analysis

Results were graphed and statistical analysis performed using GraphPad Prism Version 9 (GraphPad Software Inc., California, USA). Data are presented as the mean ± standard error of the mean (S.E.M.) for triplicate samples unless specifically mentioned. Data analysis included independent Student t-test for the comparison of two datasets or by One-Way ANOVA (Analysis of Variance) with Dunnett’s Post hoc testing with correction for multiple comparisons used to determine differences between three or more independent groups. P values of <0.05 were considered statistically significant and denoted by an asterisk in the figures.

## 3. Results

### 3.1 Aquamin inhibits LPS driven TLR4 pro-inflammatory cytokines in both primary human PBMCs and murine BMDMs

Aquamin Soluble as a nutraceutical agent was investigated for its ability to modulate pro-inflammatory cytokine levels in both human and murine macrophage primary cells. Initially we examined whether Aquamin affects the release of cytokines associated with LPS TLR4 signalling in primary human PBMCs. The effects of Aquamin treatment followed by LPS stimulation on cytokine levels in human PBMCs was measured in cell culture supernatants using ELISA assays. In the presence of LPS, Aquamin dose dependently reduced the secretion of TNF-α (Fig 1 a) and IL-6 ( Fig 1b) at 12h into the culture supernatants compared to LPS treated cells. The effects of Aquamin treatment on murine BMDMs stimulated with LPS was also investigated. BMDMs were generated by culturing murine bone marrow extracts in the presence of M-CSF (20ng/ml) for 7 days. A similar Aquamin dose dependent inhibitory effect was also observed in murine BMDMs, with TNF-α (Fig 1 c) and IL-6 ( Fig 1c) reduced at 12h post LPS stimulation (Supplemental Fig 2 b,d, 6h post LPS stimulation). This effect was also evident at the gene expression level for TNF-α and IL-6 ( Supplemental Fig 2 a,c). C-X-C Motif Chemokine Ligand 2 (CXCL2), also referred to as MIP-2 α, and C-C Motif Chemokine Ligand 2 (CCL2), also known as MCP-1, are chemokines that act as crucial signalling molecules in inflammation, attracting and orchestrating immune cells like neutrophils (CXCL2 [32]) and monocytes/macrophages (CCL2 [33]) to sites of injury or infection. Here Aquamin pretreatment of mBMDMs reduced the expression of both CXCL2 (Fig 1 e) and CCL2 (Fig 1 f) following LPS stimulation. In all cases Aquamin treatment alone did not lead to the secretion of TNF-α (Fig 1 a, b), IL-6 (Fig 1 c,d) or CXCL2 (Fig 1 e) and CCL2 (Fig 1 f) chemokine gene expression changes which were all induced upon LPS stimulation as expected. These inhibitory effects of Aquamin were not as a result of cellular toxicity, as BMDMs treated with varying doses of Aquamin (0.5, 1 and 2 mg/ml) for 12h showed no reduced cell viability compared to untreated cells (Supplemental Fig 1). These results clearly show a robust increase in the gene expression and protein secretion levels of cytokines and chemokines post LPS/TLR4 stimulation for 6h and 12h in both murine and human primary macrophages which can significantly be reduced with Aquamin treatment.

**Figure 1:**
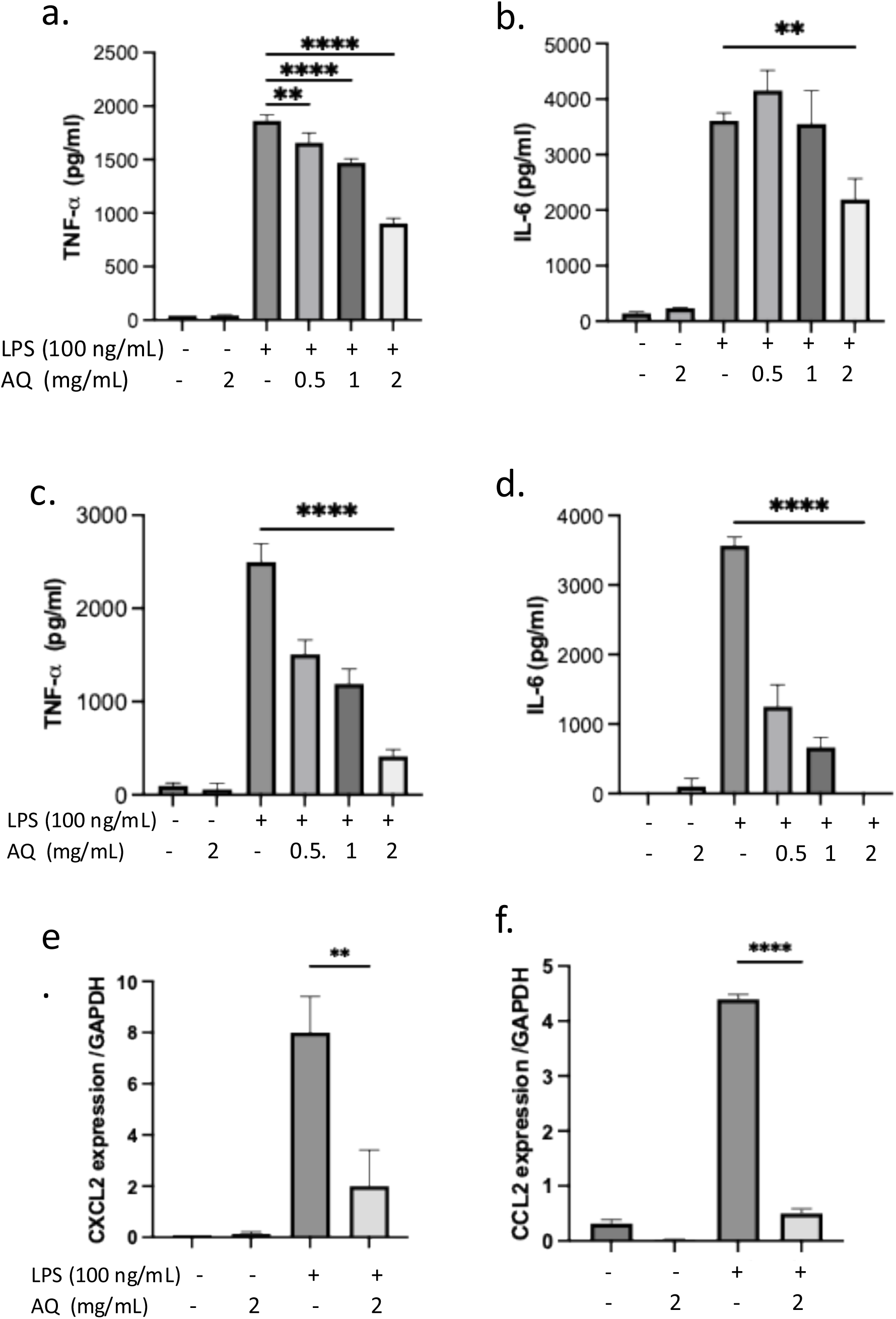
Aquamin inhibits inflammatory cytokines and chemokines downstream of LPS driven TLR 4 signaling in both human and murine macrophages. Human PBMCs were pretreated with Aquamin (0.5, 1 and 2mg/ml)) for 3 h followed by stimulation with LPS (100ng/ml) for 12h. PBMC Cell culture cytokine levels of (a) TNF-a and (b) IL-6 detected by ELISA. Murine BMDMs were pretreated with Aquamin (0.5, 1 and 2mg/ml)) for 3 h and exposed to LPS (100ng/ml) for 4 h followed by total cellular RNA extraction and 12h followed by collection of cell culture supernatants. BMDM cell culture cytokine levels of (c)TNF-a and (d) IL-6 detected by ELISA. BMDM mRNA expression levels of (e) CXCL2, (f) CCL2, normalized to GAPDH and expressed relative to untreated cells. Data presented as mean ± sem from 3 independent experiments *\*\*p< 0.01, ****p<0.0001; vs LPS alone using one way ANOVA statistical test*.

### 3.2 Aquamin inhibits LPS-induced iNOS in murine BMDM

Inducible nitric oxide synthase (iNOS) is an enzyme that catalyses the production of nitric oxide (NO) in mammalian cells. NO is an important mediator of cellular inflammation involved in pathogen killing where several immunomodulatory cytokines as well as LPS are known to upregulate iNOS and the production of NO [34]. The impact of Aquamin treatment on production of iNOS was investigated here using antibody immunofluorescence (IF) detection of iNOS in LPS stimulated BMBMs. Treatment with Aquamin alone did not increase the expression of iNOS, which was absent and similar to untreated control cells. Treatment of BMDMs with LPS in the presence of Aquamin (2 mg/ml) resulted in a decreased IF intensity of iNOS staining compared to LPS stimulation alone ( Fig 2 a, b). Representative images in Fig 2a of iNOS IF staining and DAPI nuclei counterstaining is seen in fixed BMDMs for LPS alone versus LPS and Aquamin treatment. Quantification of iNOS IF intensity normalised to nuclei DAPI staining is further shown in Fig 2b. Consistent with IF staining, iNOS protein expression levels from whole cellular lysates was quantified by western blot (Fig 2 c, d) where again the protein expression levels of iNOS was significantly reduced in Aquamin LPS treated BMDMs compared to LPS treatment alone. iNOS is a hallmark of pro-inflammatory (M1) macrophages, which produce large amounts of nitric oxide (NO) and other reactive nitrogen species (RNS) [35]. Suppression of iNOS can promote or allow for the anti-inflammatory tissue-repairing (M2) phenotype in these innate immune cells to become more prominent. Based on these results, Aquamin can inhibit the generation of an LPS mediated pro inflammatory macrophage MI phenotype here.

**Figure 2:**
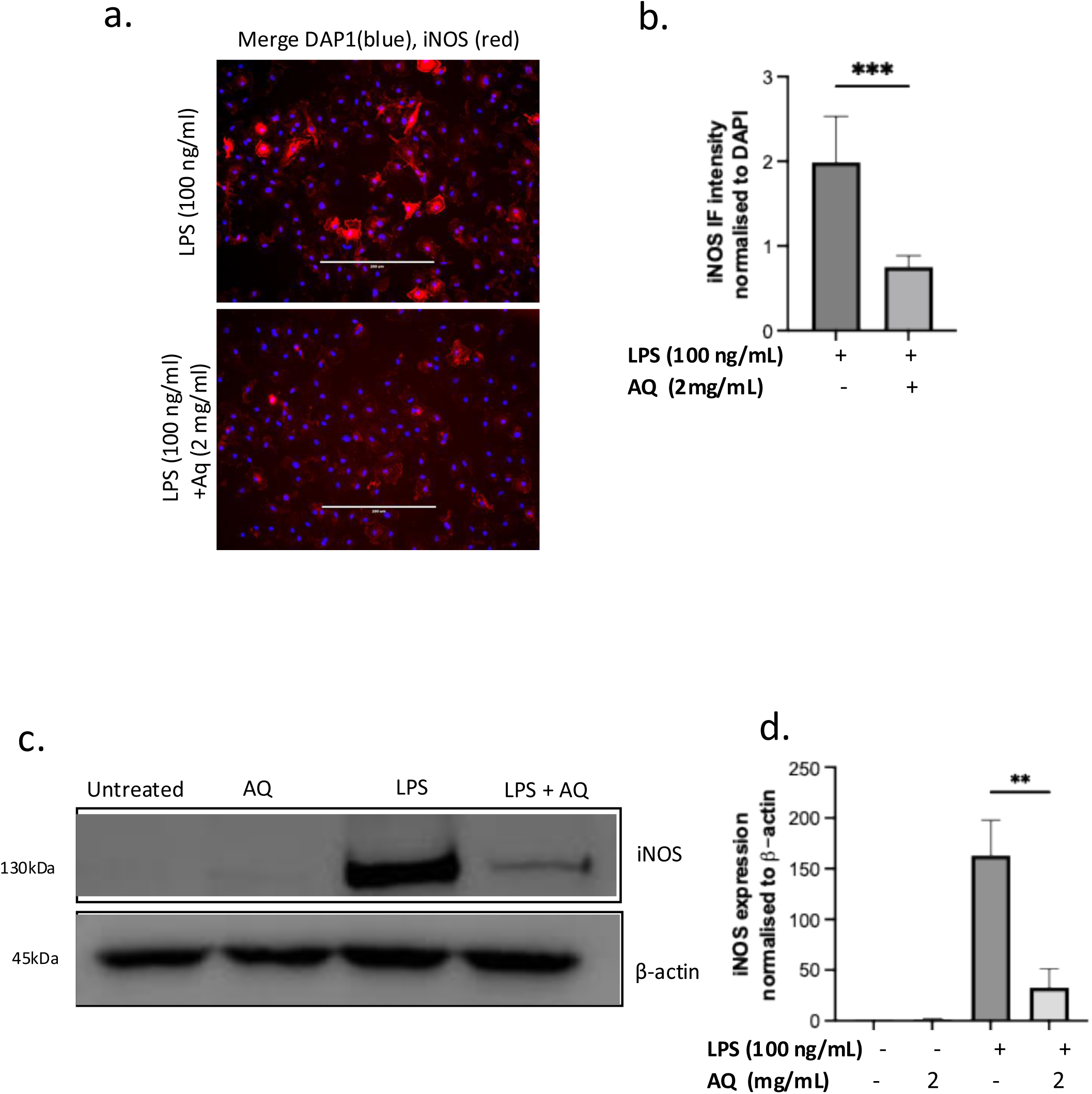
Aquamin inhibits LPS induced iNOS expression in murine BMDM. BMDMs pretreated with Aquamin (2mg/ml) for 3 h followed by stimulation with LPS (100ng/ml) for 12 h. (a) Representative immunofluorescence (IF) of iNOS protein abundance (Red), DAPI nuclear stain (blue) and actin (green) in BMDMs, scale bar (200um). (b) Bar graph of iNOS IF intensity normalized to DAPI. (c) Representative western blot protein expression of iNOS from BMDM total protein lysates. (d) Bar graph of densitometry analysis for iNOS, protein expression levels normalised to β-actin presented as fold change in protein expression relative to untreated control cells. Data presented as mean ± SEM n= 3 *\*\*p<0.01, ***p<0.001 versus LPS alone using one-way ANOVA statistical test*.

### 3.3 Aquamin inhibits TRIF gene expression downstream of TLR4 stimulation in murine BMDMs without affecting IKBα degradation and phosphorylation of p65 in murine BMDM

Similar to TNF-α and IL-6 cytokine production, LPS induction of TLR4 driven iNOS production, occurs through activation of the NF-κB transcription factor. It is well established that NF-κB activation plays a crucial role in the signalling mechanisms downstream of LPS/TLR4 in macrophage cells, driving inflammatory cytokine release [7]. This activation occurs via NF-κB p-65 subunit phosphorylation, allowing its translocation to the nucleus to initiate transcription of regulated genes. In unstimulated cells, IκBα sequesters NF-κB in the cytoplasm keeping NF-κB inactive. Upon LPS TLR4 signalling phosphorylation of IκBα occurs allowing dissociation from the NF-κB complex and IκBα targeting for proteasomal degradation [36]. To further investigate the anti-inflammatory effects of Aquamin described above, we assessed how Aquamin might impact early stage LPS induced NF-κB transcriptional activation. Interestingly, western blot protein analysis of whole BMDM cell lysates treated with Aquamin and LPS did not result in a reduction in IκBα degradation (Fig 3a b) or phosphorylation of p-65 subunit of the NF-κB complex (Fig 3 c, d) compared to LPS alone. In this setting, NF κB activation events were measured minutes after LPS stimulation indicating that Aquamin suppressive effects of TNF-α, IL-6 and iNOS are not mediated by a disruption in the early activation of NF-κB downstream of LPS TLR4 signalling. LPS TLR4 early NF-κB activation responses are dependent on MyD88 adaptor protein recruitment, whereas a delayed NF-κB response is dependent on TLR4 endosomal trafficking and TRIF adaptor protein recruitment, with TRIF also required for IRF3 transcriptional activity. As such we next sought to investigate whether Aquamin might alter the gene expression levels of both MyD88 and TRIF. Stimulation of BMDMs with LPS resulted in increased expression of both MyD88 and TRIF. However, treatment with Aquamin specifically inhibited LPS TRIF induction (Fig 3 e) without affecting LPS MyD88 gene expression levels (Fig 3 f). These findings indicate that Aquamin inhibition of TLR4 LPS signalling is not due to suppression of MyD88 dependent early NF-κB activation, but may specifically act to inhibit TRIF-dependent TLR4 endosomal signaling.

**Figure 3:**
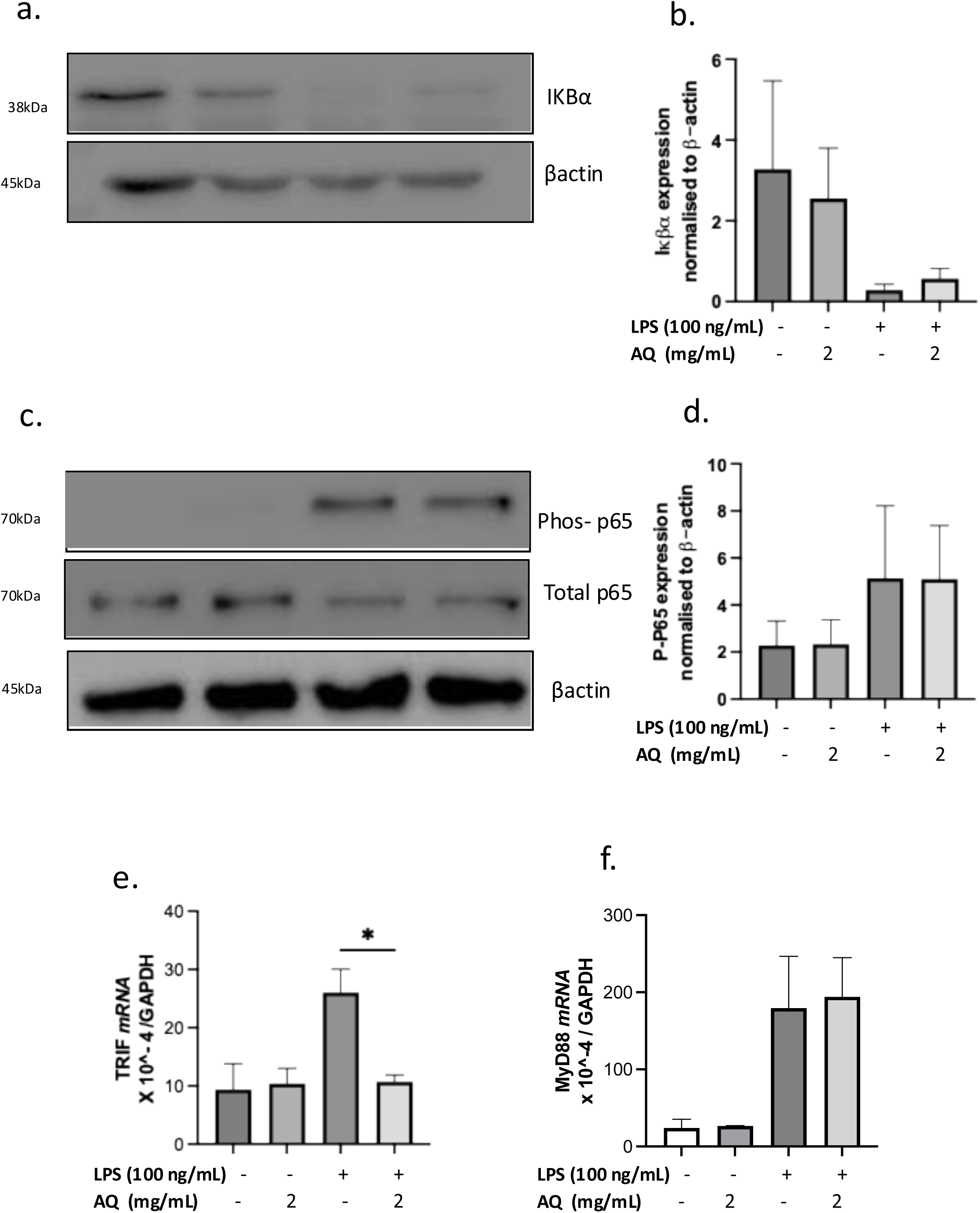
Aquamin inhibits LPS induced TRIF without affecting MyD88 gene expression, IKBα degradation and phosphorylation of p65 in murine BMDM. BMDMs pretreated with Aquamin (2mg/ml) for 3 h followed by stimulation with LPS (100ng/ml) for 15 min to detect expression of IκBα and 7 min to detect expression of p-p65 in total cell protein lysates. Representative western blot protein expression of (a) IKBα 15 min post LPS stimulation and (c) phospho p65 and total p65, 7 min post LPS stimulation. Bar graph of densitometry analysis for (b) IKBα and (d) phospho p65 protein expression levels normalised to β-actin protein loading and presented as fold change in protein expression relative to untreated control cells. BMDMs pretreated with Aquamin (2mg/ml)) for 3 h and exposed to LPS (100ng/ml) for 4h followed by total RNA extraction. (e) MyD88 and (f) TRIF gene expression normalized to GAPDH and expressed relative to untreated cells. Data presented as mean ± SEM n= 3 *\*p< 0.05 vs LPS alone using one way ANOVA statistical test*.

### 3.4 Aquamin inhibits TRIF-dependent TLR3 signalling in murine BMDM

In order to further investigate the mechanism through which Aquamin inhibits innate TLR driven responses we next investigated whether Aquamin treatment elicited any effects on other TLR family members. Unlike TLR4, which stimulates both MYD88 and TRIF dependent signalling pathways, TLR3 signalling in macrophages primarily uses the adaptor protein TRIF, without engaging MyD88 [12] [8]. To determine if Aquamin regulates TRIF dependent IRF-3 signalling, investigations using Poly (I:C) stimulation were carried out to determine if Aquamin altered the mRNA expression of the IRF-3-dependent gene, IFN-β. Using real-time PCR the mRNA expression levels of IFN-β was analysed in Poly (I:C) stimulated BMDMs pre-treated with Aquamin (2mg/ml). Aquamin reduced expression of IFN-β at the mRNA level post Poly (I:C) stimulation when compared to Poly (I:C) alone (Fig. 4 a). IFN-β expression is dependent on activation of IRF-3 through phosphorylation events, followed by IRF3 dimerization and translocation to the nucleus in macrophages after TLR3 stimulation. Therefore we next analysed the cellular localization of IRF3 expression in BMDMs under Poly (I:C) stimulation in the presence and absence of Aquamin (2mg/ml) through immunofluorescent (IF) staining. As expected, Poly (I:C) increased the nuclear localization of IRF3 as evident by nuclear DAPI counterstaining compared to untreated and Aquamin only treated cells. A reduction in the intensity of IRF3 IF staining as well as a reduction in the nuclear localization of IRF3 in Aquamin (2mg/ml) pretreated cells was evident compared to cells treated with Poly (I:C) alone ( Fig. 4 b, c). These results demonstrate that Aquamin inhibits canonical TLR3 dependent signalling responses further suggesting that Aquamin modulates TLR signalling downstream of TRIF adaptor protein pathways.

**Figure 4:**
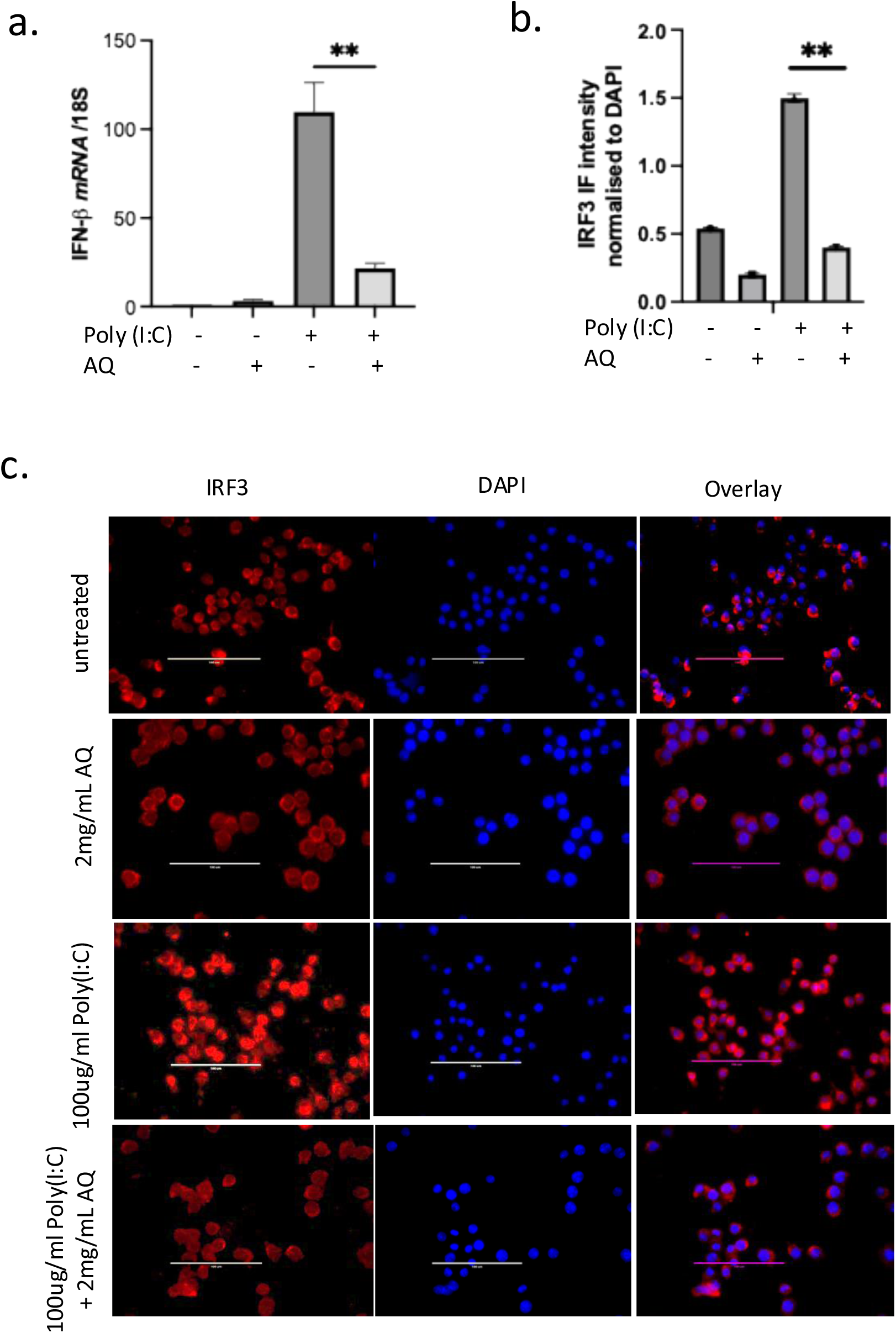
Aquamin reduced IFN-β expression and IRF3 nuclear localization in Poly (I:C) stimulated BMDM. BMDMs were pretreated with Aquamin (2mg/ml) for 3 h before stimulation with Poly(I:C) (100μg/ml) for 6 h. (a) IFN-β gene expression normalized to 18S RNA, expressed relative to untreated cells. (c) Representative immunofluorescence (IF) of IRF3 protein abundance (Red), DAPI nuclear stain (blue) and IF overlay in mBMDMs, scale bar (100 um). (b) Bar graph of IRF3 IF intensity normalized to DAPI. Data presented as mean ± sem from 3 independent experiments. *\*\*p<0.01 vs Poly (I:C) alone using one way ANOVA statistical test*.

### 3.5 Aquamin inhibits TLR4 and TLR3 but not TLR9 cytokine release in murine BMDM

Collectively, the data above demonstrates that Aquamin reduces cytokine release in human and murine innate cells under LPS and Poly (I:C) stimulation. However, these effects appear independent of MyD88 expression or activation of early downstream NF-κB mediated signalling. While TLR3 primarily signals using the adaptor protein TRIF [8], TLR9 in contrast signals exclusively using the adaptor protein Myd88 [9]. Therefore, to gain a deeper understanding of Aquamin mediated inhibition of TLR family signalling adaptor pathways we examined whether Aquamin altered BMDM responses upon stimulation with a specific agonists to TLR9 alongside TLRs 3 & 4. Primary murine BMDMs were pretreated with Aquamin for 3h and exposed to specific TLR agonists, LPS for TLR4, Poly(I:C) for TLR3, CpG DNA for TLR9 stimulation (Fig. 5) for 12h. As expected, all TLR agonists increased both IL-6 and TNF-α cytokine release from BMDMs compared to untreated cells after 12h of stimulation. While Aquamin treatment significantly reduced IL-6 and TNF-α cytokine release from LPS and Poly (I:C) stimulation, the same inhibitory effect was not evident with Aquamin upon CpG DNA stimulation (Fig 5 a,b). These data suggest that Aquamin is acting to inhibit TRIF dependent downstream signalling in response to TLR4 and TLR3 stimulation, without significantly impacting MyD88 dependent signalling pathways downstream of TLR4 and TLR9.

**Figure 5:**
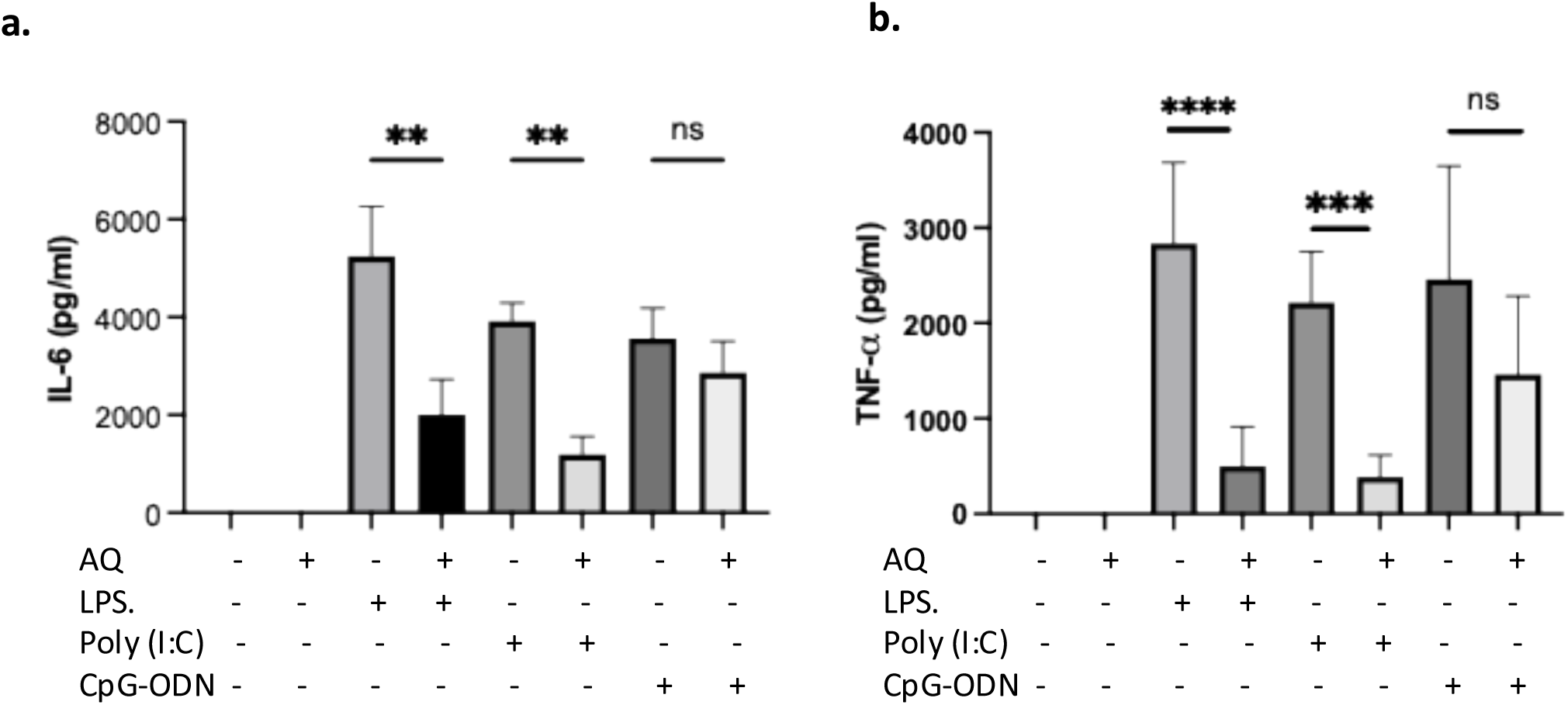
Aquamin inhibits TLR4, TLR3 stimulated release of IL-6 and TNF-α in murine BMDMs without affecting TLR9. BMDMs pretreated with Aquamin (2mg/ml) for 3 h prior to stimulation with the corresponding TLR agonists overnight, LPS (100ng/ml) for TLR4 stimulation, Poly(I:C) (100ug/ml) for TLR3 stimulation and CpG (3μM) for TLR9 stimulation. (a)IL-6 and (b) TNF-alpha cytokine levels in the cell culture supernatants by ELISA. *Data are expressed as mean ±sem from 3 independent experiments. **p<0.01, ***p<0.001 vs LPS alone using one way ANOVA statistical test*.

**Figure 6:**
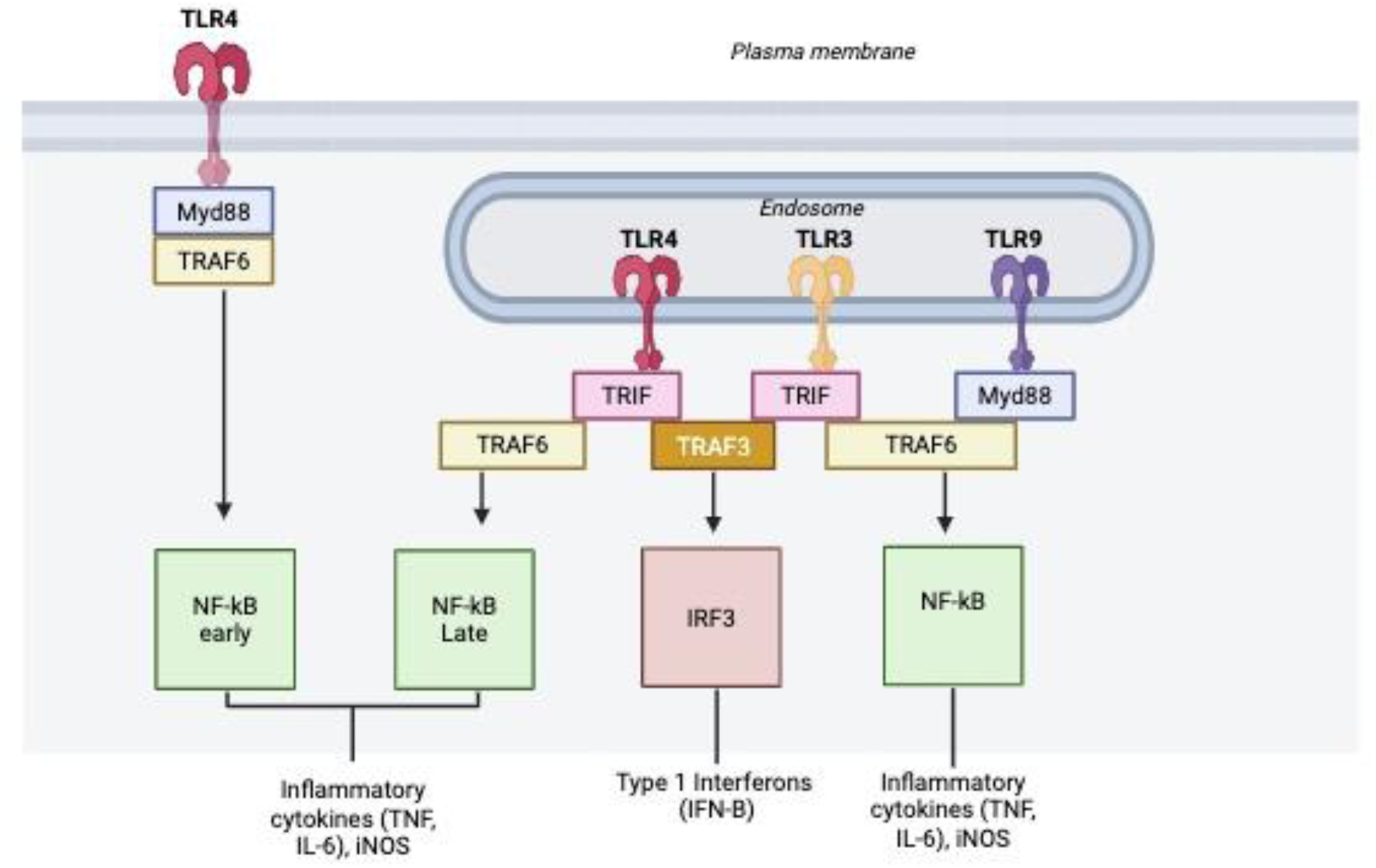
Simplified schematic of TLR 4, 3 and 9 signaling pathways. Adaptor proteins Myd88 and TRIF both act downstream of TLR4 signaling . TLR4 Myd88 initiates early NF-kB proinflammatory responses while the TLR4 TRIF pathway signaling initiates a delayed NF-kB response as well as an IRF3 type 1 interferon response. TLR3 TRIF adaptor protein signaling initiates both IRF3 and NFkB responses via TRAF3 and TRAF6 respectively. TLR9 is dependent on Myd88 signaling only to initiate NF kB responses independent of TRIF. Created with BioRender.com

## 4. Discussion

This study demonstrates that Aquamin exerts a consistent anti-inflammatory effect in primary innate immune cells by suppressing pro-inflammatory outputs downstream of TLR activation, with evidence pointing toward selective inhibition of TRIF-dependent adaptor protein signalling. In both human PBMCs and murine BMDMs, Aquamin reduced LPS-induced secretion of key inflammatory cytokines TNF-α and IL-6 (Fig 1 a-d), and in BMDMs it also suppressed LPS-driven CXCL2 and CCL2 chemokine expression (Fig 1 e,f). Importantly, these effects were not attributable to cytotoxicity (Supplemental Fig 1), indicating that Aquamin modulates inflammatory signalling rather than broadly impairing cell viability or function. Together, these findings support Aquamin as an immunomodulatory multi-mineral intervention capable of dampening macrophage-driven inflammatory programs relevant to chronic inflammatory disease states.

A notable mechanistic insight from this work is that Aquamin’s suppression of LPS-induced cytokines and iNOS (Fig 2) appears uncoupled from inhibition of early canonical NF-κB activation events (Fig 3a-d). LPS stimulation classically activates the MyD88-dependent pathway rapidly, causing IκBα degradation and phosphorylation of NF-κB p65, which drives early transcription of inflammatory mediators [10]. Here, Aquamin did not reduce IκBα degradation or p65 phosphorylation at early timepoints following LPS stimulation (Fig 3 a-d), yet downstream inflammatory outputs (TNF-α, IL-6, and iNOS) were clearly reduced at later endpoints (Fig 1,2). This suggests Aquamin does not block proximal receptor engagement or the immediate MyD88→NF-κB activation cascade. Instead, the data are more compatible with Aquamin acting on later-phase transcriptional amplification, pathway crosstalk, or adaptor regulation mechanisms that can reduce cytokine accumulation even when early NF-κB activation is preserved [36].

The adaptor protein expression data further support an Aquamin preferential modulation of a TRIF-dependent signalling response. LPS increased both Myd88 and TRIF gene expression in BMDMs, where Aquamin in the presence of LPS is shown here to selectively reduce TRIF without affecting Myd88 expression (Fig 3 e, f). TLR4 uniquely signals through both MyD88 and TRIF [8], with the selective Aquamin reduced TRIF expression over MyD88 providing a mechanistic explanation for the observed Aquamin LPS TLR4 driven immune suppressive responses in cytokine levels (Fig 1). Here the results suggests Aquamin dampens the endosomal/TRIF-dependent component of TLR4 signalling, which contributes to sustained NF-κB/MAPK activity and to IRF3-driven transcriptional gene regulation [11]. The Aquamin TRIF inhibitor response is further supported by Aquamin’s inhibitory effects observed under TLR3 stimulation, a receptor that signals primarily through TRIF [12]. Aquamin reduced Poly(I:C)-induced IFN-β mRNA expression and decreased IRF3 transcription nuclear localization, indicating suppression of TRIF–IRF3 signalling outputs (Fig 4). Collectively, the reduced TRIF expression, reduced IRF3 nuclear localization, and the reduced IFN-β induction supports the observation that Aquamin limits inflammatory responses by attenuating TRIF-dependent TLR signalling.

Adaptor molecule TLR pathway signalling selectivity was further strengthened by the agonist cytokine comparison across TLR4, TLR3, and TLR9 stimulation. Aquamin inhibited cytokine release induced by LPS (TLR4) and Poly(I:C) (TLR3) but did not significantly suppress cytokine release induced by CpG (TLR9), which signals primarily through MyD88 (Fig 5). This pattern is consistent with Aquamin preferentially targeting TRIF-dependent signalling rather than globally suppressing TLR activation. Functionally, this is an important distinction as broad inhibition of MyD88 can compromise antimicrobial defences and immune competence [37], whereas selective reduction of TRIF-driven inflammatory amplification may provide therapeutic benefit with a potentially more favourable immunological risk profile. While this study does not directly measure host defense outcomes, preservation of early NF-κB activation events and lack of effect on CpG-driven cytokines argue against nonspecific immunosuppression with Aquamin.

Aquamin also markedly reduced LPS-induced iNOS protein expression in BMDMs (Fig 2). iNOS induction is a hallmark of pro-inflammatory macrophage activation and contributes to oxidative and nitrosative stress in tissues [34]. Suppression of iNOS is therefore consistent with a shift away from an inflammatory macrophage phenotype and could be highly relevant in chronic inflammatory settings where sustained NO and reactive nitrogen species production exacerbates cellular damage by modifying proteins (S-nitrosylation) leading in the gut to potential barrier dysfunction and tissue injury [38]. Given Aquamin’s reported benefits in gut [28] [29] and liver disease models [30] reduced iNOS (Fig 2) and chemokine expression (CXCLl2 and CCL2) (Fig 1 e, f) here may help explain how Aquamin decreases immune cell recruitment and inflammatory amplification within these tissue sites supporting gut barrier integrity and limiting chronic damage.

Although the findings here point to TRIF-dependent modulation with Aquamin in TLR signalling, the precise molecular target of Aquamin remains unresolved [26]. Aquamin is a complex mixture containing calcium, magnesium, and numerous trace elements, any of which could influence inflammatory signalling [39] directly for example through kinase activity, phosphatase balance, or redox status [40] or indirectly through membrane dynamics and receptor trafficking. One plausible mechanism consistent with the data is that Aquamin influences TLR4 endosomal trafficking or the formation and stability of TRIF-associated signalling complexes, which would preferentially dampen TRIF-dependent outputs while leaving early MyD88-driven events largely intact. Alternatively, Aquamin may regulate transcriptional control of TRIF itself or alter post-translational events required for IRF3 activation and nuclear translocation. Distinguishing among these possibilities will require targeted follow-up experiments assessing TRIF protein levels, IRF3 phosphorylation/dimerization, endosomal TLR4 localization, and measuring downstream signalling intermediates (e.g., TBK1/IKKε activation).

In this study we recognise the mechanistic conclusions are based on adaptor mRNA regulation and functional readouts where a direct assessment of TRIF protein abundance and activity would strengthen the model. Second, early NF-κB signalling was evaluated using IκBα degradation and p65 phosphorylation; however, NF-κB transcriptional outcomes depend on additional co-factor recruitment, and MAPK-driven transcriptional synergy [36], which were not measured. Furthermore studies investigating cell-type-specific profiling such as macrophage polarization assays would clarify which populations are most responsive and whether Aquamin promotes broader anti-inflammatory programming [35].

In conclusion, Aquamin robustly suppresses TLR-driven inflammatory mediator production in primary human and murine innate immune cells and appears to do so through preferential inhibition of TRIF-dependent signalling rather than the early MyD88-dependent NF-κB activation cascade. This TRIF selectivity—supported by inhibition of TLR3 responses and lack of inhibition of TLR9 responses—provides a mechanistic framework that aligns with Aquamin’s reported protective effects in inflammatory disease contexts[28, 29]. Future work defining the proximal molecular target(s), confirming TRIF/IRF3 pathway inhibition at the protein and signalling-complex level, and validating these effects in relevant in vivo models will be essential to establish Aquamin’s therapeutic potential as a nutraceutical strategy for controlling pathological inflammation while preserving core innate immune responsiveness.

