## Supplemental Data for "Aquamin a marine derived multi mineral attenuates toll like receptor mediated inflammatory responses in macrophages"

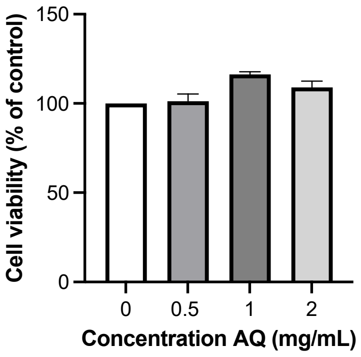

**Supplemental Figure 1 :** Cell viability measured using resazurin assay in murine BMDMs treated with Aquamin (0.5, 1 2 mg/ml)  overnight. Data is presented as percentage of control untreated cells.

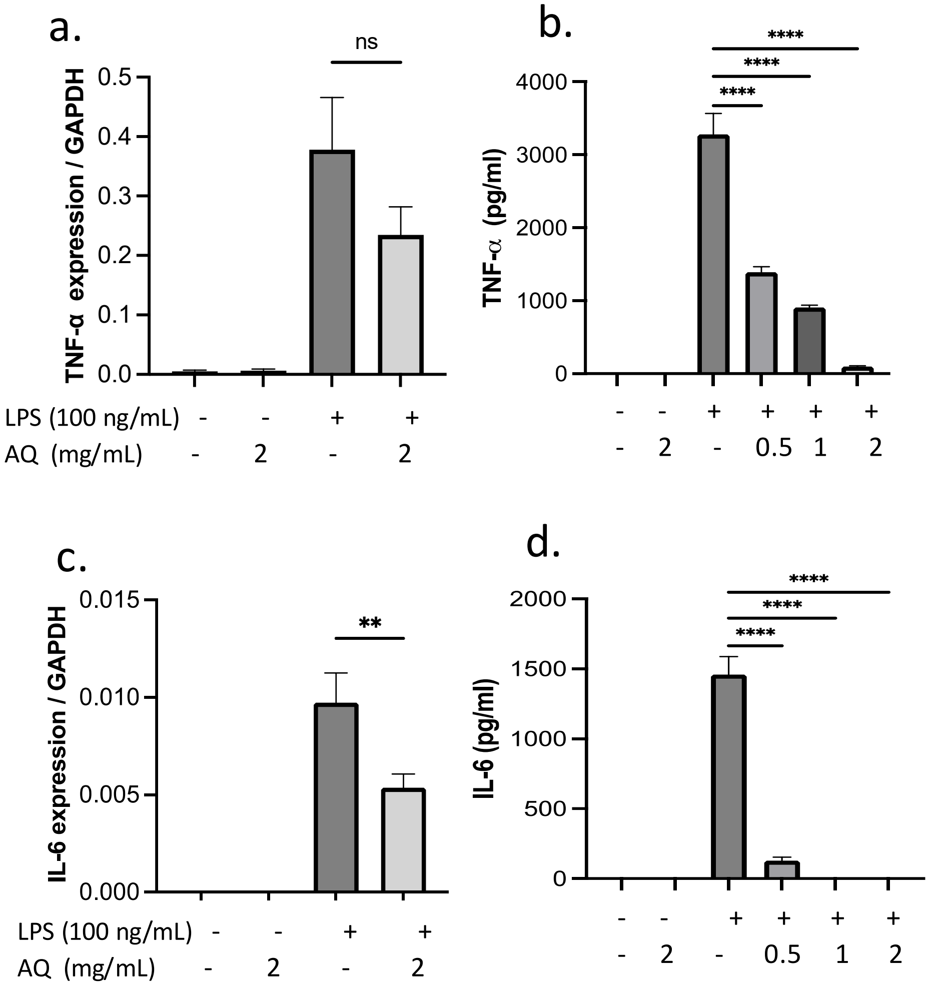

**Supplemental Figure 2 : Aquamin inhibits TNF-α and IL-6 downstream of LPS driven TLR 4 signaling in murine BMDMs.** BMDMs were pretreated with Aquamin (0.5, 1 and 2mg/ml)) for 3 h and exposed to LPS (100ng/ml) for 4 h followed by total cellular RNA extraction or 6h followed by collection of cell culture supernatants. The mRNA expression levels of (a) TNF-a, (c) IL-6, presented as mean ± sem from 3 independent experiments normalized to GAPDH and expressed relative to untreated cells. Cytokine concentrations of TNF-a (b) and IL-6 (d) detected in the cell culture supernatants by ELISA **p< 0.01, ****p<0.0001; vs LPS alone using one way ANOVA statistical test.

**Supplemental Table 1 – SYBR® Green primer sequence**

| Gene Name |  | Gene Sequence |
|---|---|---|
| GAPDH | Forward | 5'- AAC AGC AAC TCC CAC TCT TC- 3' |
|  | Reverse | 5'- CCT GTT GCT GTA GCC GTA TT- 3' |
| TNF-alpha | Forward | 5'-CTA CCT TGT TGC CTC CTC TTT-3' |
| | Reverse | 5'- GAG CAG AGG TTC AGT GAT GTA G-3' |
| IL-6 | Forward | 5'- GTC TGT AGC TCA TTC TGC TCT G-3' |
| | Reverse | 5'- GAA GGC AAC TGG ATG GAA GT-3' |
| TRIF | Forward | 5'- GGA CCT CAG CCT CTC ATT ATT C-3' |
|  | Reverse | 5'-AGG TTC CCT TCC TCC ACT AT-3' |
| MyD88 | Forward | 5'- GTA TCC TGC GGT TCA TCA CTA T-3" |
| | Reverse | 5'- GAA CTC TTC CAC TCA GCT ATC C-3' |
| CXCL2 | Forward | 5'-CCA TTG CCC AGA TGT TGT TAT G -3' |
| | Reverse | 5' GCC ATC CGA CTG CAT CTA TT-3' |
| CCL2 | Forward | 5'- GAA GGA ATG GGT CCA GAC ATA C -3' |
| | Reverse | 5'- TCA CAC TGG TCA CTC CTA CA -3' |
| IFN-beta | Forward | 5'-GGA AAG ATT GAC GTG GGA GAT-3' |
| | Reverse | 5'-CAG GCG TAG CTG TTG TAC TT-3' |
| 18S RNA | Forward | 5'-CTG AGA AAC GGC TAC CAC ATC-3' |
| | Reverse | 5'-GCC TCG AAA GAG TCC TGT ATT G-3' |

| **Supplemental Table 1. Composition of Aquamin®** | | | | | |
|---|---|---|---|---|---|
| **Element** | Amount | Element | Amount | Element | Amount |
| **Carbon** | 255,000 ppm | Indium | <0.001 ppm | Samarium | 0.035 ppm |
| **Aluminum** | 22.6 ppm | †Iodine | 0.41 ppm | Scandium | 1.209 ppm |
| **Antimony** | <0.5 ppm | Iridium | 0.002 ppm | Selenium | <0.5 ppm |
| **Arsenic** | 0.101 ppm | Iron | 80.1 ppm | Silicon | 9.44 ppm |
| **Barium** | 4.06 ppm | Lanthanum | 0.899 ppm | Silver | 1.28 ppm |
| **Beryllium** | 1.60 ppm | Lead | 0.019 ppm | Sodium | 2,055 ppm |
| **Bismuth** | <0.5 ppm | Lithium | <0.5 ppm | Strontium | 968 ppm |
| **Boron** | 13.8 ppm | Lutetium | 0.012 ppm | Sulfur | 1,144 ppm |
| **Cadmium** | 0.268 ppm | Magnesium | 1.05% | Tantalum | 0.015 ppm |
| **Calcium** | 12.10% | Manganese | 17.03 ppm | Tellurium | 6.67 ppm |
| **Cerium** | 0.225 ppm | Mercury | 0.004 ppm | Terbium | 0.008 ppm |
| **Cesium** | <0.001 ppm | Molybdenum | 0.773 ppm | Thallium | <0.5 ppm |
| **†Chloride** | 1,852 ppm | Neodymium | 0.143 ppm | Thorium | 3.09 ppm |
| **Chromium** | 1.27 ppm | Nickel | 0.605 ppm | Thulium | 0.005 ppm |
| **Cobalt** | 1.51 ppm | Niobium | 7.77 ppm | Tin | 0.215 ppm |
| **Copper** | 2.01 ppm | Osmium | <0.001 ppm | Titanium | 2.72 ppm |
| **Dysprosium** | 0.046 ppm | Palladium | 0.143 ppm | Tungsten | <0.5 ppm |
| **Erbium** | 0.033 ppm | Phosphorus | 20.4 ppm | Vanadium | <0.5 ppm |
| **Europium** | 0.012 ppm | Platinum | 0.008 ppm | Ytterbium | 0.030 ppm |
| **†Fluoride** | 2.099 ppm | Potassium | 83.2 ppm | Yttrium | 2.71 ppm |
| **Gadolinium** | 0.039 ppm | Praseodymium | 0.031 ppm | Zinc | 1.76 ppm |
| **Gallium** | 0.037 ppm | Rhenium | 0.001 ppm | Zirconium | 3.48 ppm |
| **Germanium** | <0.001 ppm | Rhodium | 0.147 ppm | | |
| **Gold** | <0.5 ppm | Rubidium | 0.011 ppm | | |
| **Hafnium** | 0.074 ppm | Ruthenium | 0.473 ppm | | |
| **Holmium** | 0.010 ppm | | | | |

The trace mineral composition of Aquamin® TG was determined by an independent laboratory (Advanced Laboratories, Inc., Salt Lake City) for Marigot Limited (Ireland) in 2018.

Individual trace element composition was determined by Inductively Coupled Plasma Optical Emission Spectroscopy (ICPOES) except for carbon (determined by ASTM D-1552), chloride and iodine (determined by Titration), and fluoride (determined by AOAC 939.11).

**Supplemental Table 3 Microbiological testing screen**

| Total viable count | 5,000cfu/g max. |
|---|---|
| **Yeast & Moulds** | 100cfu/g max. |
| **E. Coli** | Absent in 1g |
| **Coliforms** | Absent in 1g |
| **Enterobacteriaceae** | Absent in 1g |
| **Staphylococcus aureus** | Absent in 1g |
| **Salmonella** | Absent in 25g |
